# Nonlinear Neural Patterns Are Revealed In High Frequency fNIRS Analysis

**DOI:** 10.1101/2022.08.03.502588

**Authors:** Ameer Ghouse, Diego Candia-Rivera, Gaetano Valenza

## Abstract

Vasomotor tone has a direct implication in oxygen transport to neural tissue, and its dynamics are known to be under constant control from feedback loops with visceral signals, such as sympathovagal interactions. Functional Near Infrared Spectroscopy (fNIRS) offers a nuanced measure of hemoglobin concentration that also comprises high frequencies, though most fNIRS literature studies traditional frequency ranges of hemodynamics (< 0.2 Hz). Linear theory of the hemodynamic response function supports this low frequency band, but we hypothesize that nonlinear effects elicited from the complex system sustaining vasomotor tone presents itself in higher frequencies. To characterize these effects, we investigate how plausible modulation of autoregulatory effects impact aforementioned high frequency components of fNIRS through simulations of mechanistic hemodynamic models. Then, we compare representational similarities between fast (0.2 Hz to 0.6 Hz) and slow (< 0.2 Hz) wave fNIRS to demonstrate that representations acquired through nonlinear analysis are distinct between the frequency bands, whereas when using linear time-domain analysis they are not. Furthermore, by comparing topoplots of significant detectors using nonlinear random vector correlation methods (distance correlation), we demonstrate through a 2nd level group analysis that the median concentrations acquired by fNIRS are independent when analyzing the nonlinearity of their dynamics in their fast and slow component, while they are dependent when utilizing linear time-domain analysis. This study not only provides motivation for researchers to also include higher frequency components in their analysis, but also provides motivation to explore nonlinear effects, e.g. topological entropy. The results of this study motivate future research to explore the nonlinear autoregulatory impacts of regional blood flow and hemoglobin concentrations.

**Author summary:** Conventionally, hemodynamic response from induced neural metabolic demand is studied as a slow signal, i.e < 0.2 Hz. Though this may be justified in linear analysis of hemodynamics, vascular mechanics nonlinearly transform the neural metabolic demand to hemodynamic response, where a nonlinear spectral profile may show higher frequency responses. Higher frequency ranges may give insight into local vascular dynamics, particularly their reflection of autoregulatory phenomena, hypothesized to be controlled by sympathovagal feedback loops, thus opening a new avenue for studying brain-body interactions. Functional near infrared spectroscopy (fNIRS) offers a method with high temporal resolution (10 Hz) for observing these effects in hemoglobin concentrations. In this study, we utilize stochastic dynamical simulations of plausible autoregulatory phenomena and an open fNIRS dataset to study differences of fast and slow wave neurovascular representations. We demonstrate that, while linear time-domain analysis provides similar representations of fast and slow wave activity, representations derived from nonlinear methods are not. Furthermore, we show how stress tasks, which may elicit autonomic activity, further desynchronizes nonlinear activity between fast and slow wave signals compared to a non-stress inducing task, demonstrating unique high frequency neurovascular phenomena that is mediated by stress processing.

## Introduction

Autonomic system activity contains complex dynamics derived from feedback loops in baroreflex sensors and nonlinear sympathovagal interactions [1]. Indeed, many physiological systems demand full characterization via nonlinearity and complexity analysis [2–4]. Blood flow has been speculated to have induced responses to autoregulatory feedback. Autoregulatory feedback and its interaction with the nonlinear autonomic activity has been given great debate in literature. [5–12]. Nonetheless, hemodynamic models [13] contain parameters that correspond to vasomotor tone and resistance [14, 15] where there is evidence of its modulation by autonomic activity [11, 12], justifying the reflection of how these parameters’ dynamics affect components in the brain, particularly oxygen transport whose crucial role as an electron acceptor in aerobic respiration sustains neural activity [16]. Indeed, biomarkers of sympathovagal interactions with the brain, such as brain-heart interplay, have also provided evidence that visceral signals such as heart beat control precede cortical activations [17, 18], which similarly puts into demand the role of vasoreactivity in cortical oscillations.

On a neurophysiological level, autonomic activity and stress has shown to be strongly intertwined [19] in a similar way to how autonomic system is connected to emotions [20, 21]. In fact, stress regulation can similarly be seen as emotional regulation [22]. It has been shown that indeed there is an information exchange between the brain and the visceral signals when studying the brain-heart axis as a proxy for sympathovagal-cortical interactions during stress [23]. Though, the stress and emotion arousal effects may diverge in the neurophysiology [21, 24, 25], having a comprehensive regional autonomic system activity response measure with the brain interactions provides an axis that can aid in studying the brain-body interactions during psychological emotional disorders using an all-in-one method.

Functional near infrared spectroscopy (fNIRS) provides a method to measure brain tissue hemoglobin concentration as a proxy for neural activity [26]. Slow hemodynamic response signals (< 0.2 Hz) are theorized to be elicited by neurogenic stimuli [13, 27, 28], and are well approximated by linear analysis when time intervals between stimuli are long [28], thus fNIRS analysis commonly focus on the hemodynamic frequency band, and high frequency signal is filtered out due to the assumption that it corresponds to systemic physiological or instrumentation noise [29]. Furthermore, cut-off frequencies are conservatively low (< 0.2 Hz) to avoid overlap with frequency components in vascular dynamics that are independent with brain function (such as cardiac oscillations). However, the system is indeed nonlinear [13–15], and may potentially be impacted by the aforementioned nonlinear autonomic system control [11, 12, 14, 15]. Thus, linear time invariant proposals of hemodynamic spectral activity may underrepresent the true underlying phenomenon, motivating investigation into the nonlinear effects in higher frequencies (> 0.2 Hz) Implementation of nonlinear signal analysis often lift observed time series to a manifold in higher dimensional phase space via delay-coordinate map as specified in Takens’ theorem [30]. Entropy is one of the measures used for characterizations of time series in such spaces, and Sample entropy (SampEn) [31] may be measured in phase space to assess irregularity based on the correlation integral that describes the probability that a state is in a partition of a smooth manifold.

Though traditional analysis of fNIRS focuses on frequencies less than 0.2 Hz so as to avoid overlap with frequency components in vascular dynamics that are independent to brain function [32], cognitive stress from mental arithmetic has been assessed using fNIRS heart rate derived signal in Hakimi et al [33]. However, these high frequency components were primarily used as an extra indicator for classifying stress levels rather than assessing induced spatial activity changes. Furthermore, there exists little justification for substantial effects of cardiac rhythms on local cortical areas. Most evidence points to anatomical disruptions associated with local cardiac pulsatility changes [34]. Regardless, there is difficulty in motivating conclusions on regional effects instead of systemic body effects from variations in pulsatility. Past research in Ghouse et al [35] assessed entropy estimates of fNIRS signals that not only contained the traditional hemodynamic and heart rate frequencies jointly band passed, it also contained an upper bound of 0.6 Hz rather than the lower suggested cutoff frequencies of fNIRS preprocessing [32], and demonstrated complementary areas of activity when compared to neural correlates observed using linear analysis of fNIRS. This raises the question of what additional information may have been added by this extra range of frequency in hemodynamic fNIRS signal to reveal the complementary areas of activity.

Thus, the aim of this study is to assess differences between neural representations of fNIRS in traditional slow wave (< 0.2 Hz) and proposed fast wave (0.2 Hz to 0.6 Hz) frequencies. First, simulations of plausible modulations of autoregulatory feedback on blood flow control in the hemodynamic model [13] is explored to motivate fast wave nonlinearity analysis. Then experimental data is exploited, using both a mental arithmetic paradigm and a control motor imagery paradigm [36] to study the additional information high frequency components may provide when a subject is put under cognitive stress. We perform a representational similarity analysis (RSA) [37, 38], i.e. a multivariate pattern analysis (MVPA) that has shown power in assessing whether modalities (such as a theoretical model, behavioral data, empirical data, etc) contain similar representations for cognitive states–here we apply it to comparing fast and slow wave fNIRS modalities. Experimental data is obtained from a public database as provided by Shin et al. [39]. Furthermore, we assessed multivariate correlation at each distance correlation to assess whether the topoplots median differences between tasks are correlated in fast and slow wave analysis across the group. We hypothesized that fast wave activity would add significant complementary regions, particularly in nonlinear analysis instead of linear analysis, according with the results obtained in [35]. Furthermore, we expected mental arithmetic topoplots of activity to be less correlated between fast and slow wave fNIRS activity as per the hypothesis of autoregulation and autonomic system interactions affecting blood flow regionally, with nonlinearities present in frequency ranges outside of traditional bands. Particularly, mental arithmetic is hypothesized to be less correlated between fast and slow wave fNIRS when compared to the correlations during a motor imagery task [36].

## Materials and methods

### 0.1 Simulations

For initial validation of plausible effects of autoregulation dynamics on hemoglobin, we first simulate a mechanistic balloon model of hemodynamic activity [13, 40]. First, some flow inducing signal (s), i.e. the response of the vascular system to a neural metabolic demand, is linearly described for the sustenance of incoming blood flow (*f_in_*).

Particularly:

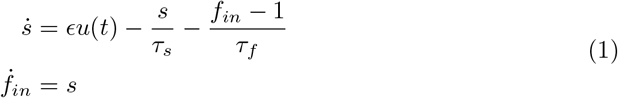

u is the control signal which is the neural signal generating the flow. *ϵ* is the efficacy by which the neural signal can sustain a flow-inducing signal, i.e. a response that may dilate vessels to increase blood flow into. This is related to the vascular resistance (r), whose dynamics are 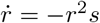 [13]. *τ_s_* is the time constant for this flow-inducing signal and *τ_f_* is the time constant describing the effects of autoregulatory feedback from blood flow. Considering *τ_f_* comprises information on autoregulatory effects of blood flow (the time constant for returning back to baseline), this is the parameter we later model to assess its nonlinear effects on hemoglobin concentrations.

From the sustaining blood flow generating signal, a so-called “balloon model” describes the dynamics of the volume of a blood vessel (the balloon), and the permeation of hemoglobin inside and outside the vessel [41]. Explicitly, the rate of decay of blood volume is related to blood flowing in and blood flowing out of a vessel:

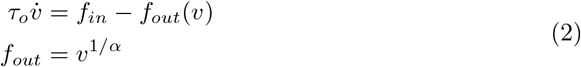

*α* describes the capacity at which a balloon can expel water, having been distended by the surge of inflow. *τ_o_* is the time constant which governs the rate of change of the volume, which is similarly intertwined with the rate of change of deoxyhemoglobin in the venous compartment (q):

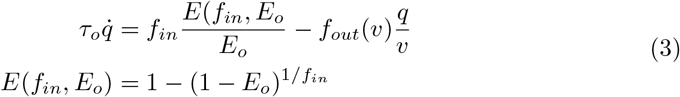

The function E describes the oxygen extraction coefficient, or how efficacy of the tissue in extracting oxygen from the incoming blood, while *E_o_* is the resting oxygen extraction coefficient. In Cui et al [40], an extension was proposed to relate the blood volume and total hemoglobin concentration as:

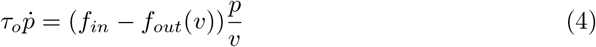

Then, oxyhemoglobin is merely *o* = *p* – *q*.

We reiterate the observation in eq. 1 that the parameter *τ_f_* is particularly related to the autoregulatory feedback. Exactly how its value relates to the true autoregulatory changes is little understood, though literature states that responses to autoregulatory changes occur over a period of 1-2 minutes (or less than 0.02 Hz) [5]. To maintain this expected periodicity, while allowing random deviations due to uncertain dynamics, we designed a 2nd order stochastic dynamical model whose power spectral density on average has a peak at the frequency of hypothesized autoregulatory changes.

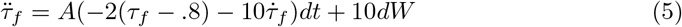

The steady state value of *τ_f_* is at 0.8, while the steady state of its change is 0. A is a parameter that modulates how fast it returns to steady state, and the diffusion term is random variations outside the potential normal oscillatory behavior of *τ_f_*. When there is no autoregulatory activity, it quickly returns to steady state, with A=10. When there is autoregulatory activity, it is more free to change with A=.1. We integrate the stochastic differential equations with the Euler-Maruyama method [42].

We numerically simulated 100 *τ_f_* time courses to illustrate expected spectral properties of the stochastic dynamics. Furthermore, we simulated a block design experiment, each block being 40 seconds long with a minute long rest, where 5 stimulus blocks would either induce *τ_f_* modulations (i.e. where A = 0.1) or 5 stimulus blocks would not induce the modulations. The final simulated hemoglobin concentration time courses are corrupted with a signal-to-noise (SNR) ratio of 0 dB.

### 0.2 Experiment Design

A publicly available dataset was used to obtain fNIRS signals with the desired experimental protocol for this study, as reported in [39]. In summary, the experiment recruited twenty-nine healthy subjects (aged 28.5 ± 3.7), fifteen of which were female and fourteen male. Three trials were performed with ten repetitions of mental arithmetic and baseline events, and three trials were performed with ten repetitions of right hand and left hand motor imagery for each subject. We note that the motor imagery tasks and mental arithmetic tasks were done independently, not concurrently. Thirty-six fNIRS series were acquired for each subject with 10 Hz sampling rate.

The experiment design had sixty seconds of resting state to start data acquisition from a subject, after which an instruction was shown on the screen indicating which task was to be performed–either an arithmetic problem, a “-” for a baseline, or a left or right arrow for motor imagery. The subject performed the indicated task for ten seconds, with a subsequent fifteen second resting phase before the next instruction. After twenty repetitions of these instructions and tasks (ten second repetition per task), a sixty second rest was performed. A total of three trials were performed, for a total of thirty repetitions per event.

### 0.3 fNIRS signals

Thirty six channels of optical densities (OD) were resolved from source detector pairs comprising 760nm and 850nm wavelengths covering the frontal, lateral parietal and posterior cortical regions as seen in 1a. The modified Beer Lambert law was used to convert the ODs to deoxyhemoglobin (Hb) and oxyhemoglobin (HbO) [43].

Figure 2 illustrates the preprocessing pipeline. After applying the modified Beer-Lambert law, band-pass frequency filters were applied to extract traditional hemodynamic bands (< 0.2 Hz) [32, 43] or the proposed increased hemodynamic band (0.2 Hz to 0.6 Hz). A wavelet filtering approach using a Daubechies 5 wavelet, nine level decomposition was used to further reduce instrumentation noise such as movement in the oxy- and deoxyhemoglobin signals [44]. Detrending and prewhitening with an AR(1) model was then performed to remove temporally structured noise in the signal [45]. The signals were separated into epochs, with each channel at each activity block being referenced to the mean of the previous 5s. Total hemoglobin was computed as the addition of both Hb and HbO. For all three signals, entropies and mean estimates were extracted then passed into the 1st and 2nd level analysis. Before second level analysis, the results were spatially smoothed to increase the sensitivity of random effects from detector locations [46]. A Gaussian smoothing kernel was used as seen in fig. 1b.

**Fig 1.**
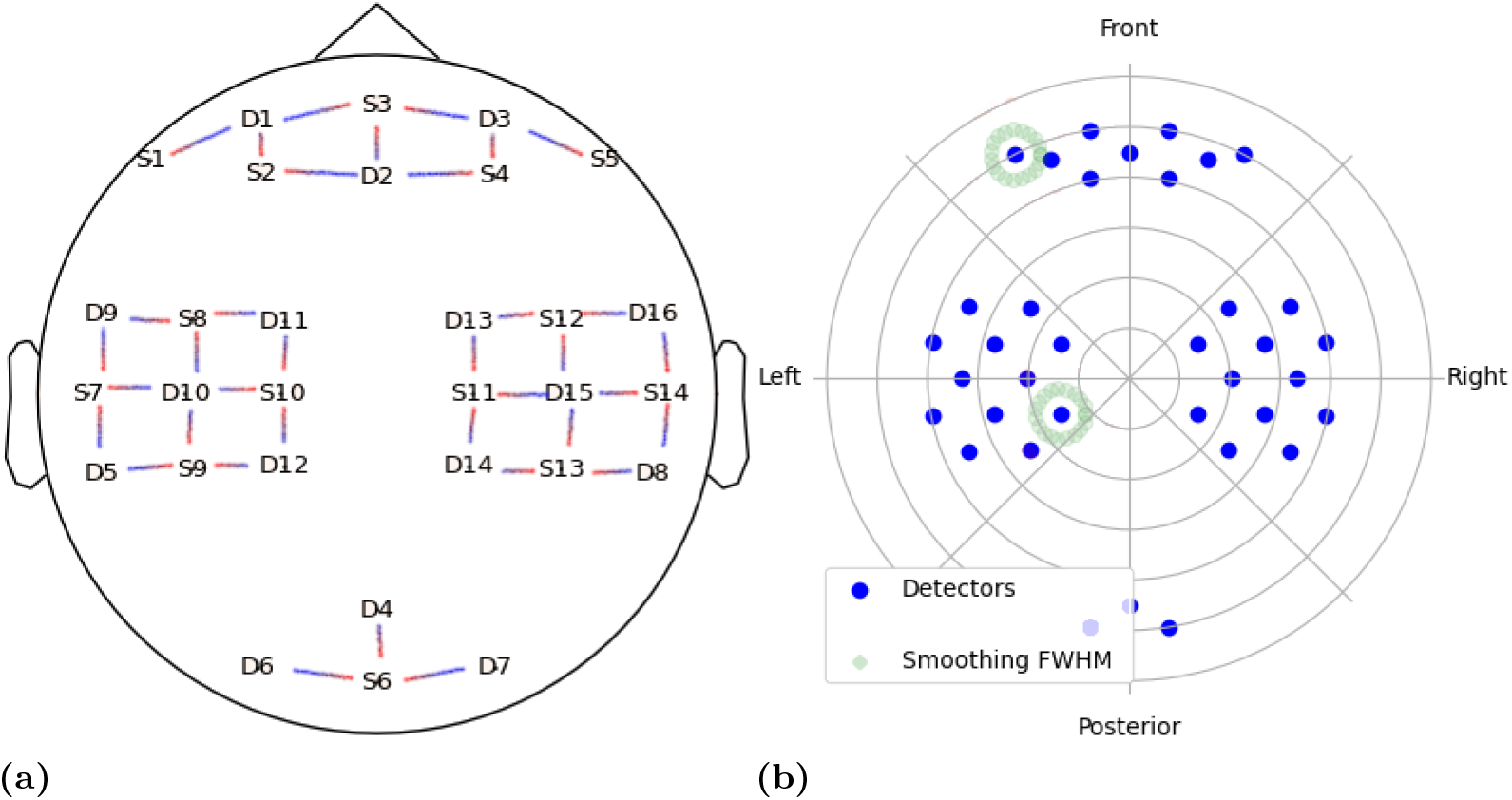
Illustration of spatial analysis of fNIRS signals. (a) demonstrates the position of the optodes used to derive the 36 fNIRS channels. (b) shows an example of the full-width-half-max of the smoothing Gaussian kernel prior to second level group analysis in the color green

**Fig 2.**
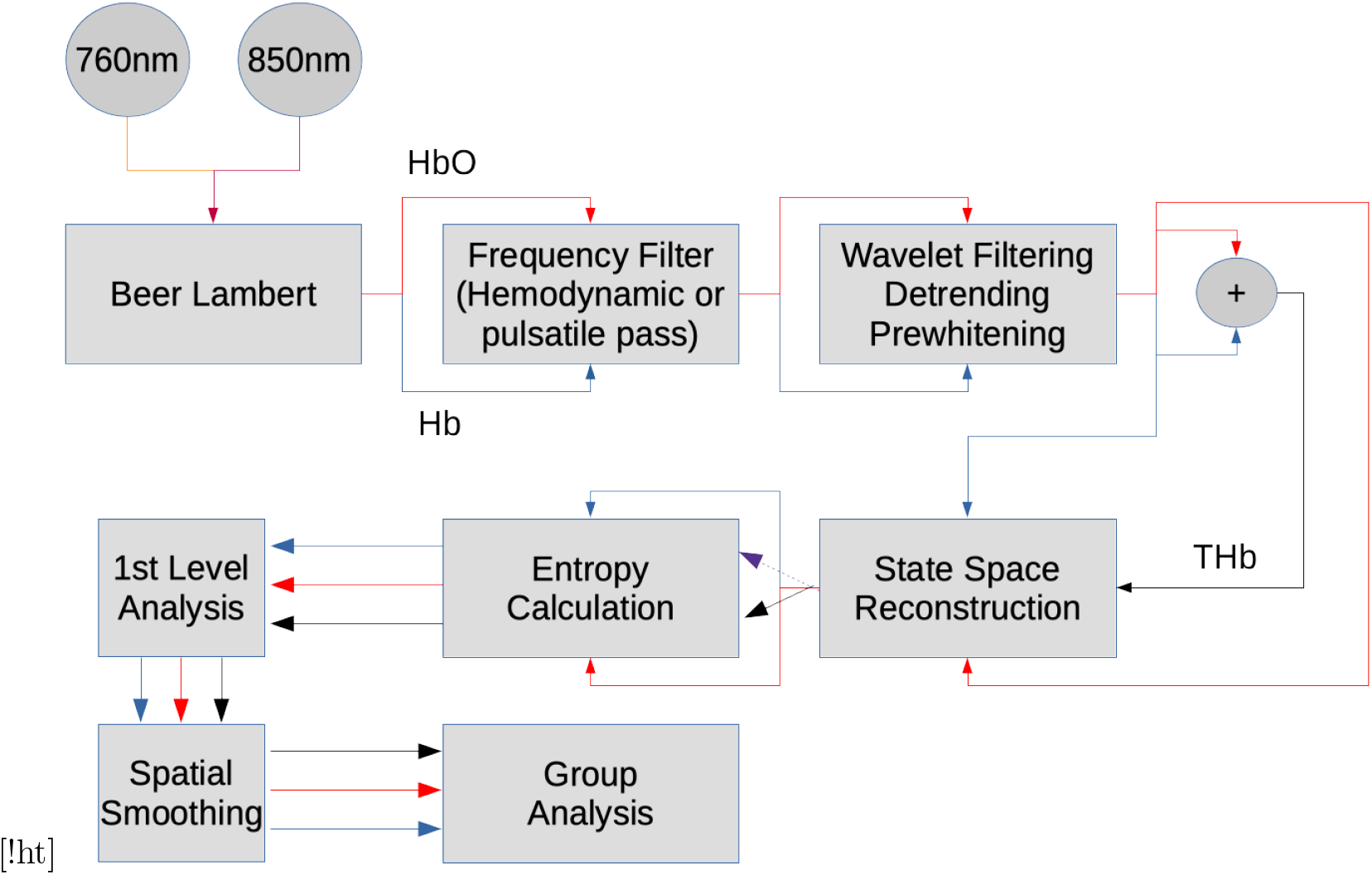
Analysis pipeline used for each fNIRS signal in the dataset

### 0.4 Entropy Analysis

A delay-time *τ* and embedding dimension *m* are needed to reconstruct manifolds using delay-coordinates. *τ* was selected as the first zero of the autocorrelation, while m was found using the false nearest neighbors approach [47].

For calculating SampEn, radius *R* = 0.2 × *σ_x_* was used as the threshold to determine whether states were neighbors, where *σ_x_* is the standard deviation of the fNIRS time series [48]. The particular equation describing how to calculate SampEn is:

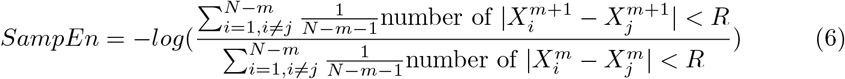

X in this equation is a state space reconstructed using a time series. The superscript denotes the embedding dimension while the subscript denotes the state index, for which there exists N states.

Beer-Lambert law derives both a time series for HbO and Hb. From adding these two time series, a third time series, total hemoglobin (THb) can be computed. Thus, an embedding of a manifold in a state space given by lag coordinates can be reconstructed from each of these three time series.

### 0.5 Representational Similarity Analysis

Each detector’s data per subject had a shape of N repetitions × N concentrations, where concentrations correspond to hemoglobin concentrations. To derive dissimilarity between mental arithmetic and baseline representations (which are random vectors), the distance correlation was used [49, 50]. Briefly, given random vector X and random vector Y with dimensionality ℝ^*p*^ and ℝ^*q*^ respectively, alongside their respective characteristic functions (i.e. Fourier transform of their probability density functions) *ϕ_x_*(*t*) = *E*(*e^itX^*) and *ϕ_y_*(*t*) = *E*(*e^itY^*), distance correlation tests independence by integrating the distance between the random vectors along with a weighting function 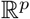 where 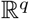 is a function corresponding to half the surface area of a unit sphere in the given dimensionality *d.* This results in the following statistic:

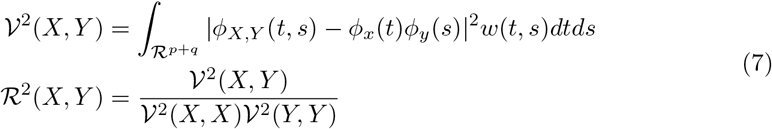

As the dimensionality of the random vectors, p and q, go to infinity, the statistic approaches a student t distribution and can be approximated as one for performing hypothesis tests [51]. Given *a_ij_* = |*x_i_* – *x_j_*| and *b_ij_* = |*y_i_* – *y_j_*|, where i and j are the ith and jth observations of x or y, the sample covariance can be estimated as:

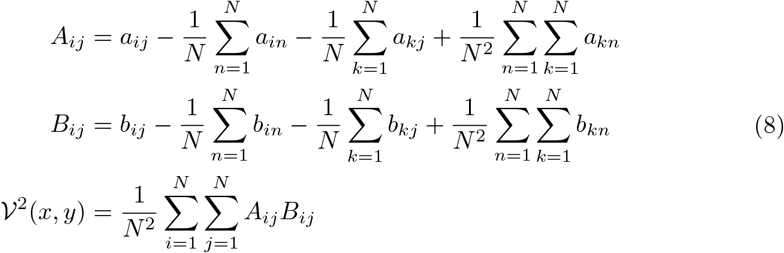

Other correlation measures such as Pearson or Spearman are based on random variables rather than random vectors, thus are not applicable. Due to there being 4 experimental conditions (baseline, mental arithmetic, left and right hand imagery), the resulting representational dissimilarity matrix (RDM) is a 4 × 4 matrix.

Representational dissimilarity outputs were obtained for each detector location for each measure (entropies or mean value). Extracting the upper triangle, we have two 29 matrices for slow and fast wave fNIRS respectively to perform statistical analysis upon.

### 0.6 Fast and Slow Wave Spatial Analysis

Using distance correlation, we can similarly compare the spatial distribution of the difference of median values between the concentrations over the mental stress inducing trials and the motor imagery trials, treating the spatial array of difference of median values of each concentration for a subject as a random vector for either fast or slow wave fNIRS. That means each subject has a vector N Concentrations × N detectors. Such an analysis will allow a comparison to determine whether fast and slow waves spatial fNIRS arrays are dependent, or in other words the null hypothesis that they are independent can be rejected. We would expect that the slow and fast wave analysis would become more independent when there is a task inducing changes in autoregulatory activity. Beyond the distance correlation, we apply Wilcoxon paired analysis to compare, at a detector level, which medians are significantly different in the group of subjects for each concentration.

### 0.7 Statistical Analysis

Group analysis for the representational similarity analysis is performed to compare RDMs between fast and slow wave fNIRS using a Friedman test. A Bonferroni correction was used (i.e. multiplying the p-value by thirty-six detector center comparisons) to correct for multiple comparisons. A p-value < 0.05 is considered significant after the multiple comparisons correction.

For median analysis, rainclouds are generated from standardized Z (i.e. zero mean and unit variance) transformations of the measures in order to be able to compare shapes of distributions between fast and slow wave analysis. We are obliged this processing step as the scales of fast and slow wave analysis, especially in mean analysis, are not the same thus demanding such a transformation for shape comparison. For comparing distributions of concentrations analyzed from fast and slow wave analysis for either SampEn or mean, we applied a Kolmogorov-Smirnov test, correcting for multiple comparisons (4 tasks and 3 concentrations; *α* = 0.05/12 = 0.0042). Then, a paired sample Wilcoxon signed rank test was performed on the difference of median value of for each concentration between the activities of interest (either mental arithmetic vs baseline, or right hand vs left hand motor imagery). A detector was retained for further analysis if any of the 3 concentrations returned significant with a false alarm rate *α* = 0.05/6 = 0.008, where the value 6 was the correction for the multiple comparisons over 3 concentration values and 2 modalities (fast or slow wave analysis). For the remaining detectors, the distance correlation is performed between fast and slow wave fNIRS.

## Results

Results are from simulations, as well as group statistics of representational similarity analysis, and median analysis of topoplots. Standard fNIRS analysis refers to time averaged value during an event, while fNIRS nonlinear analysis refers to SampEn during the time duration of the event.

### 0.8 Simulations

Fig. 3 demonstrates the results of realizations of the hypothesized autoregulatory feedback changes as according to eq. 5. The median peak frequency was at 0.02 Hz, corresponding to periodicity of 50 seconds, with periodicity ranges from 12 seconds to 100 seconds. Fig. 4 illustrates the effects of that these autoregulatory effects may have on the hemodynamics, where HbO is the oxyhemoglobin and HbR is the deoxyhemoglobin. Over the 100 realizations of the block design experiment, the mean entropy of the hemodynamic response in the high frequency (0.2 Hz to 0.6 Hz) regime without autoregulation modulations was at 0.212 (±0.015) bits compared to 0.163 (±0.011) bits when the modulations occur; through a paired Wilcoxon signed rank test, the SampEns were found to be significantly different (*p* ≪ 0.001). On the other hand, the power of this high frequency band without autoregulation modulation was found to be 0.0368 (±0.00441) as compared to 0.0372 (±0.00385) during autoregulation modulation; the power bands were not significantly different according to the Wilcoxon signed rank test (*p* = 0.46).

**Fig 3.**
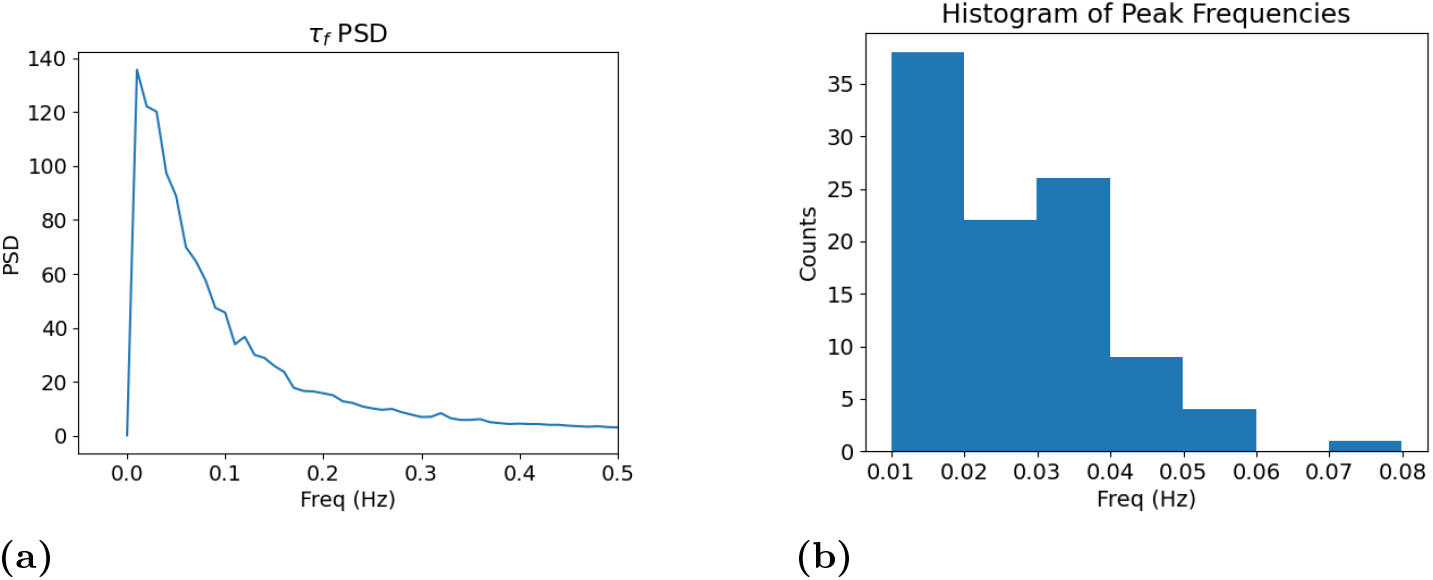
Spectral analysis of realizations of *τ_f_* simulated from the stochastic differential equation in eq. 5. a) Represents the average PSD of *τ_f_* as estimated by the Welch method over 100 realizations. b) on the other hand is a histogram of the peak frequency of the PSD of *τ_f_* over the 100 realizations

**Fig 4.**
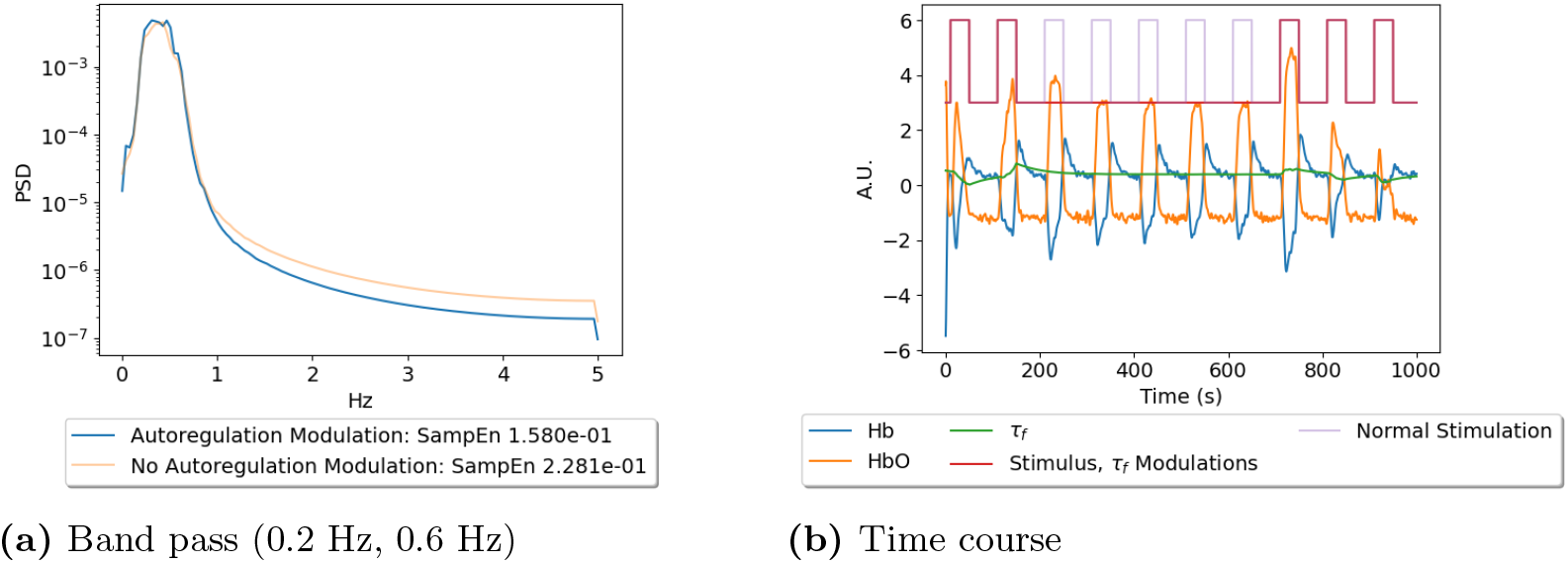
(a) demonstrates a single realization of a band passed oxyhemoglobin power spectrum density when either autoregulatory activity is occurring with a task, or when there is no autoregulatory activity with a task. (b) displays a representative simulation of the signals in the model and the events that influence them (whether a task induces autoregulatory behavior or not).

### 0.9 Group Statistics of Representational Similarity Analysis

Results from the Friedman analysis comparing each the upper triangle of the RDM can be seen in fig. 5, for mean and SampEn respectively. For the mean value, no detector has significantly different representations between slow and fast wave fNIRS, whereas each quadrant of the sensor space on the cortex contained significantly different detectors for SampEn. Generally, SampEn contained higher Friedman test scores than the mean analysis.

**Fig 5.**
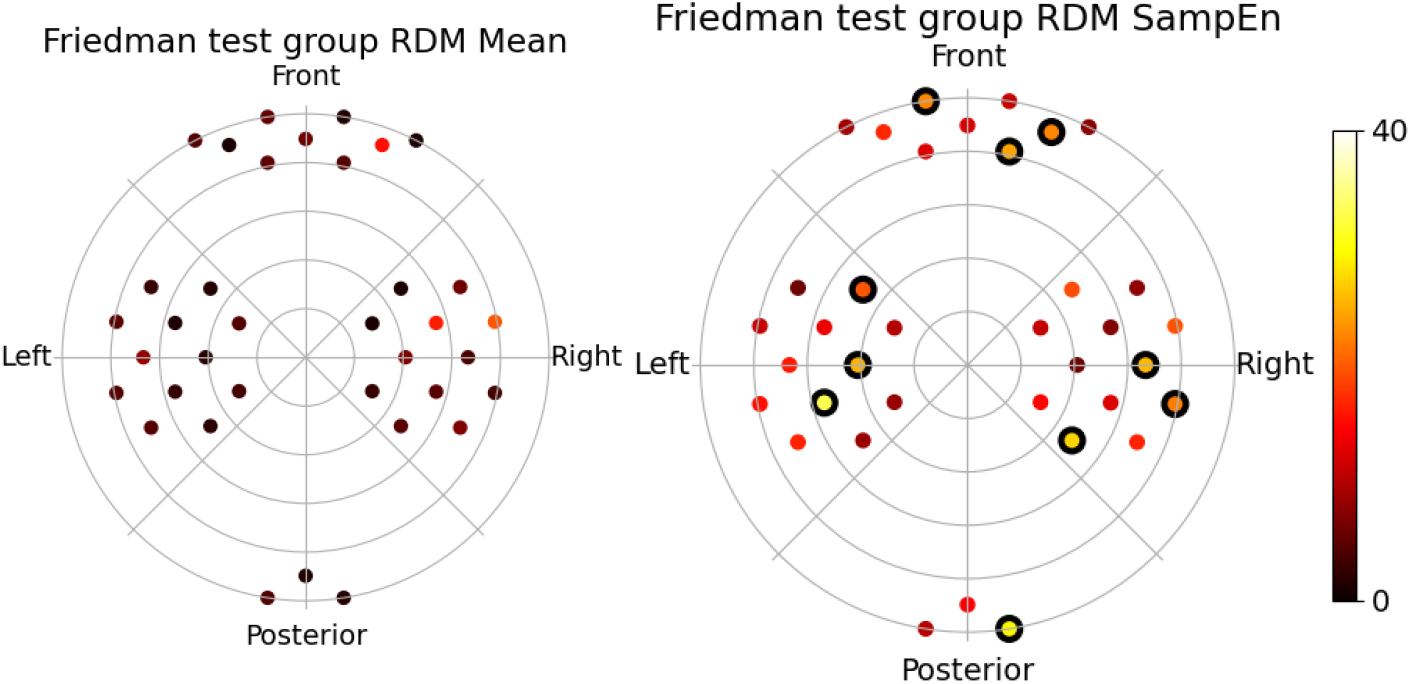
Results of a group level Friedman test comparison between the upper triangle of the RDM for fast (0.2 Hz to 0.6 Hz) and slow wave (< 0.2 Hz) fNIRS on the group level, where significance indicates at least one element in the upper triangle had a significantly different median between fast and slow wave fNIRS. Channels that are significant are marked by a thick outline on the marker.

### 0.10 Group Statistics of Median Analysis

Group level median analysis results for fast and slow wave fNIRS can be seen in the raincloud fig. 6 and topoplot fig. 7 for the comparison between mental arithmetic (MA) and baseline (BL), or right (RH) or left hand (LH) motor imagery tasks.

**Fig 6.**
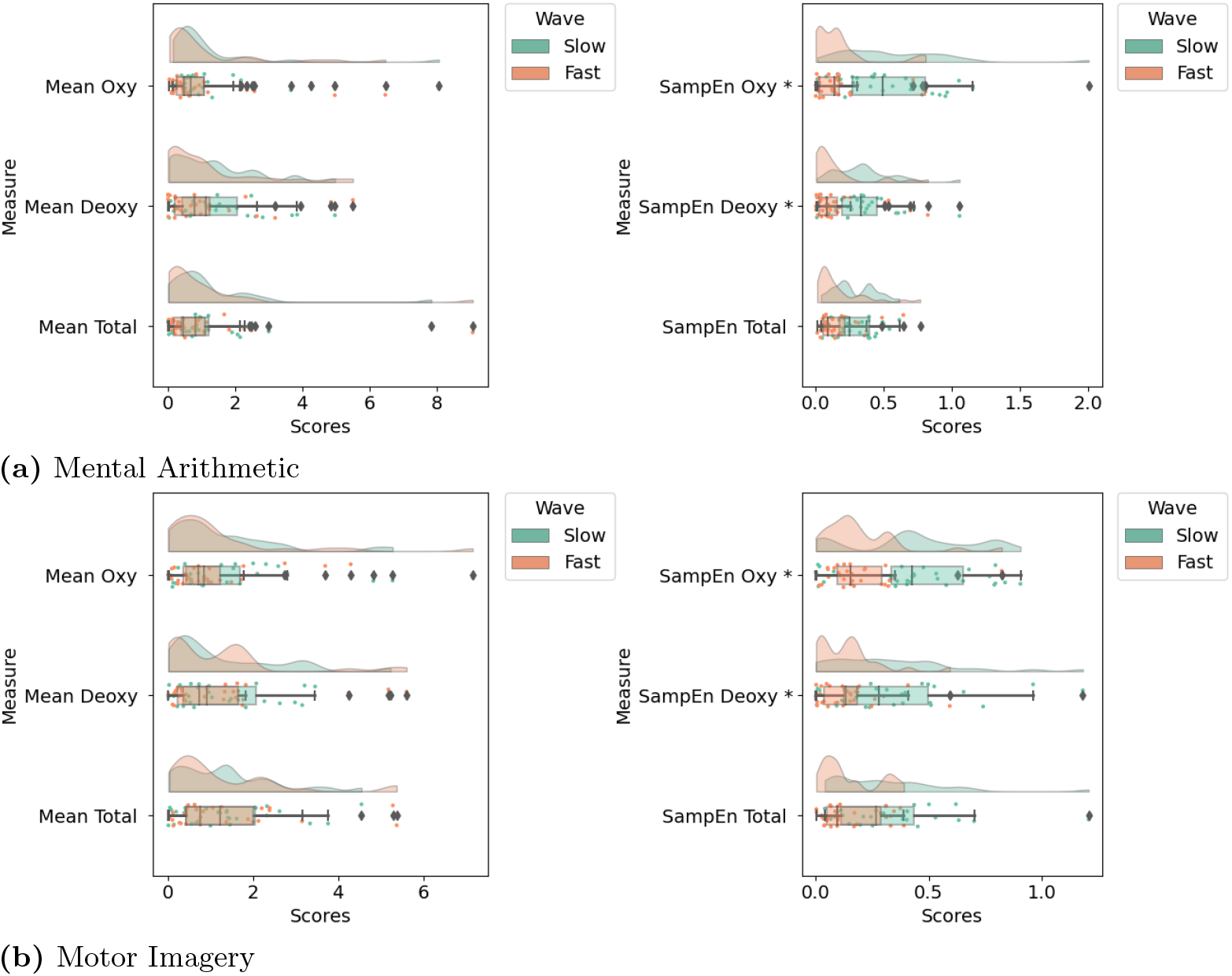
Rainclouds illustrating the standardized Z transformed group level distributions of the absolute value of the difference between medians for mental arithmetic (a) and motor imagery (b) for either fast wave (0.2 Hz to 0.6 Hz) or slow wave (0 Hz to 0.2 Hz) fNIRS signals. The “*” represents significant difference between the fast and slow wave distribution for the given measure.

**Fig 7.**
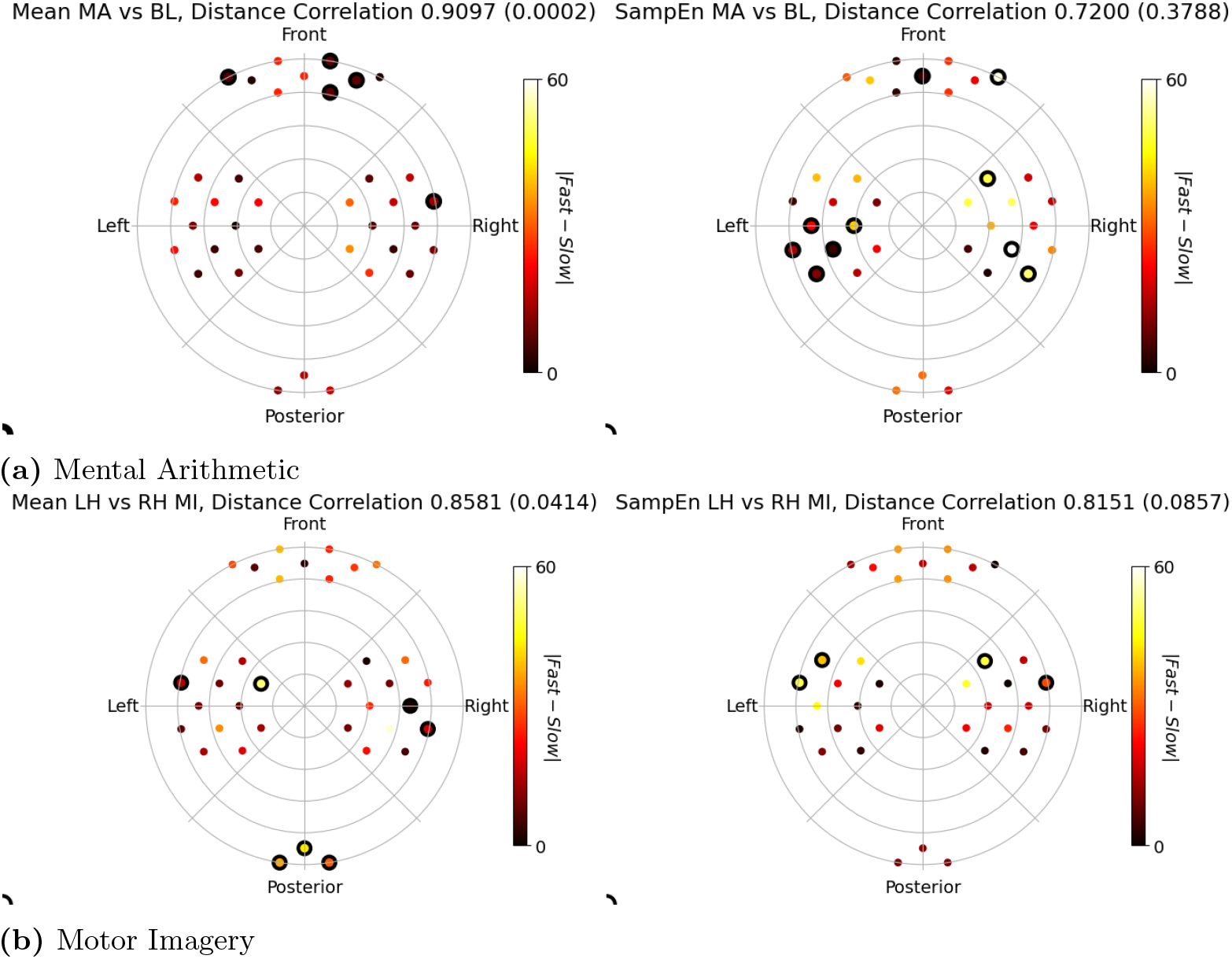
Results from comparing fast wave fNIRS (0.2 Hz to 0.6 Hz) and traditional slow wave fNIRS (0 Hz to 0.2 Hz). Effect size displayed in the plots are the absolute value of the difference between the medians at the group level between fast and slow wave measures (either mean in the left column or sample entropy in the right column). Group analysis Wilcoxon signed rank tests were used on the group level to assess whether a detector center has a significantly different median for two tasks, where surviving detectors are demarcated by a thick marker outline. (a) displays mental arithmetic vs baseline results, while (b) displays results comparing left vs right hand motor imagery. The colors in the plots represent which modality (fast or slow wave) was significant for the given measure. In the title of each subplot, the distance correlation among the surviving detectors between fast and slow wave topoplots is provided, with the p-value in parenthesis.

From the raincloud plots in fig. 6, distributions for mean results consistently appear to be overlapped when comparing slow and fast wave different in medians, whereas SampEn demonstrates a flatter distribution for slow wave analysis as compared to fast wave analysis. Motor imagery also presents a flatter distribution in SampEn when using slow wave analysis compared to fast wave analysis, however this greater variance is similarly observed in mean value analysis. Performing a Kolmogorov-Smirnov two sample test, we found that the distribution of fast and slow wave analysis was significantly different between SampEn in oxy and deoxyhemoglobin, while in no mean analysis were they significantly different.

For mental arithmetic and baseline Wilcoxon paired analysis, mean estimates show significant detectors only in slow wave fNIRS, with a cluster in the right lateral cortical areas and detectors bilaterally in the frontal cortex. SampEn mainly determined significant detectors in the frontal right cortex and left parietal cortex in the slow wave analysis, while fast wave analysis complemented information in the right parietal cortical areas.

Left and right hand motor imagery Wilcoxon paired analysis demonstrated activity bilaterally in the parietal cortical areas with both SampEn and mean estimates. Both mean and SampEn demonstrated slow activity predominantly in the right parietal cortex, while fast activity was found in the left lateral parietal cortex. Mean also contained significant detectors in the medial left parietal cortex in slow wave analysis and in the occipital regions for fast wave analysis.

Distance correlation for mean analysis in either task comparison were statistically significant, demonstrating that the topoplot results were not ever independent events. SampEn on the other hand were never significant for rejecting the hypothesis of independence, with mental arithmetic demonstrating even lower correlation than motor imagery.

## Discussion

In this study, representations of neural activity were compared between fast wave (0.2 Hz to 0.6 Hz) fNIRS and the standard slow wave (< 0.2 Hz) fNIRS for both linear measures of mean value and nonlinear measures derived from entropies in phase space. The fNIRS signals were taken from a publicly available dataset, described in [39]. 29 subjects were analyzed with one set of trials performing mental arithmetic with baseline tasks, and another set of trials performing left and right hand motor imagery tasks.

It was hypothesized that the activity which induces cognitive stress would provide more variation in areas of cortical activation than motor imagery when comparing slow and fast wave fNIRS when applying nonlinear analysis. This hypothesis was particularly motivated by literature suggesting linear characteristics of hemodynamics are primarily slow (on the order a 5 seconds for the hemodynamic response to reach its peak [28]), while being aware that hemodynamic models are nonlinear in nature [13, 27, 41]. The nonlinear models signals are sustained by a variety of factors that may be induced by visceral affects that affect vasomotor tone, for example [11–15], which speculatively may be initiated be sympathovagal interactions [5–12].

To begin motivating nonlinear spectral analysis of the hemodynamic signal given our hypothesis that such hidden variables that affect vascular resistivity induce nonlinear effects in hemodynamic signals fNIRS observes [11–15], we simulated potential autoregulation modulations caused by vasomotor property dynamics, according to eqs. 1 and 5. Fig. 3 demonstrated that our theoretical model for modulations of vasomotor tone follows a similar spectral profile to what we expect real world vasomotor tone variations to be at with frequency less than 0.02 Hz [5]. Furthermore, we demonstrated that these autoregulation variations for flow signals that propagate hemoglobin concentration dynamics, as seen in fig. 4 decrease the entropy in the observed signal even with an SNR of 0 dB for the simulated observed signal while no appreciable differences can be gleaned for a direct comparison of the PSD (a linear analysis). This gives confidence that, provided a proper power of statistics, such variations induced by autoregulatory dynamics can potentially be seen in real world data, particularly when analyzed with nonlinear methods.

We attempted to see whether any element in the RDMs over the tasks of baseline, mental arithmetic and motor imagery were the same between fast and slow wave fNIRS, using a Friedman test, as seen in fig. 5. For mean value, none of the combination of tasks were significantly different, thus we had to accept the null hypothesis that the medians were the same. This is to be expected as the mean corresponds to a DC value of the signal, and a high pass filter merely attenuates that DC value from slow to fast wave fNIRs, thus not affecting their representations. SampEn contained significant areas of difference between fast and slow wave fNIRS for detectors in all 4 quadrants of the sensor space covering cortex, and generally had a higher Friedman score across the cortex compared to mean analysis. It is not evident exactly how the filters affect the regularity of state space, however Borges et al show that high pass filtering results in lower mean entropy but higher variance than low-pass filtering [52]. This effect may correspond to the significantly different SampEn representational similarity across subjects seen.

From assessing the standardized Z-transformed distributions in 6, we illustrated that fast wave analysis contain distributions that are less flat than the slow wave analysis distributions, while in mean analysis the two distributions appeared to have the same shapes. This further confirmed the different distribution of representations of neural signal between fast or slow wave analysis. Using the group analysis of median concentrations with the Wilcoxon test, as seen in fig. 7, we finely parsed through which regions on the cortical surface may contain concentrations that are different between mental arithmetic or baseline in either slow or fast wave fNIRS. Mean was particularly only sensitive to detecting differences in the slow bands, while SampEn found difference in activities when either using slow or fast wave fNIRS. Both entropy and mean estimates uniquely found information in the frontal right cortex in slow wave fNIRS, agreeing with the literature [53, 54]. Parietal information was consistent across fast wave fNIRS for all measures for SampEn, though laterality was different between fast and slow wave fNIRS; left parietal cortex seemed particularly involved in slow wave analysis while fast wave analysis found more lateral information in the right hemisphere. Left parietal activity matches activity that corresponds to processing of numbers while right parietal activity corresponds to decisions involving numbers [55]. Thus, it is not as if one is more correct than the other, rather perhaps the processing of numbers (i.e. the evaluation of a mathematical expression) induces more slow changes, while the decisions we make on numbers (i.e. should we subtract or add) induces more irregular waves in fast frequencies. One would expect with mean analysis, similar to what was found in the RSA analysis, that fast wave and slow wave would provide the same cortical regions of information; we only observe slow wave activity. This could be due to the fact that the high pass filter was performed on the entire trial time series rather than only at the block of activity, thus artifacts may arise due to the nonstationarity of a signal, i.e. the full time series statistics may not be the same as the statistics of a time series at a window.

Regarding motor imagery, it appeared that significant detectors changed laterality in SampEn when considering which frequency of band was of focus, where fast wave analysis was more sensitive to left hemisphere activity while right hemisphere activity was more sensitive to slow waves. A similar phenomenon was seen in the mean analysis, too, through there was an additional medial left hemispheric detector was observed for slow waves. We expect activity to correspond to the somatosensory cortex, which is located on the central parietal cortical regions, where hands of the cortical homunculus being located between the lateral and medial portions of the cortex [56]. We may have discovered its bilateral effects when considering the analysis of both frequency bands using SampEn. This bilateral effect was found using only slow wave analysis using just the mean analysis. This may suggest that to describe the complementary areas of activity that entropy provides, the full frequency profile of potential hemodynamic activity must be taken into consideration.

From the distance correlation of the difference of median maps as seen in 7, all the mean maps were significantly correlated as expected. The cortical representations as reflected in median difference maps between tasks should be similar if we scale the signal merely by a DC value as discussed previously. However, with our hypothesis, we expected mental arithmetic to not be significantly correlated in entropy analysis while motor imagery should be significant. We instead observe that both of them are not significant, albeit mental arithmetic vs baseline has lower correlation than motor imagery distinction. Speculatively, this may imply the nonlinear irregular effects induced by vasomotor mechanical dynamics are always occurring, though the degree to which it affects the resulting neural signal is different according to the task. Indeed, it is known that a large portion of cerebral vascular resistance is from vasomotor control in arterioles [57–60]. The nonlinear transform of the constant dynamics of vasomotor control to the hemoglobin concentrations may always be present, indeed, yet their scale is modulated by a stress task such as mental arithmetic, making their high frequency contributions to the hemoglobin concentrations more irregular in the case of mental arithmetic. As an additional sanity check, to ensure representations are unique when uniqueness is known, we assess a comparison between the traditional frequency ranges of hemodynamics with the cardiac pulsatile signal which exists in fNIRS [61]. Our hypothesis here would be that these should always be unique between fast and slow wave fNIRS when assessing the irregularity of the signal, as the cardiac component is thought in fNIRS instrumentation to be a purely systemic effect, thus not sensitive to the actual process sustaining cognitive activities [29]. Indeed, recreating fig. 7 using “fast wave” signal on the band (0.8 Hz, 3 Hz) for the cardiac component, we see that the representations are indeed not significantly correlated in the nonlinear estimates of irregularity with sample entropy, seen in fig. 8. And the detectors that are significant with fast wave using the cardiac frequency has minimal overlap with the significant detectors seen in fig. 7.

**Fig 8.**
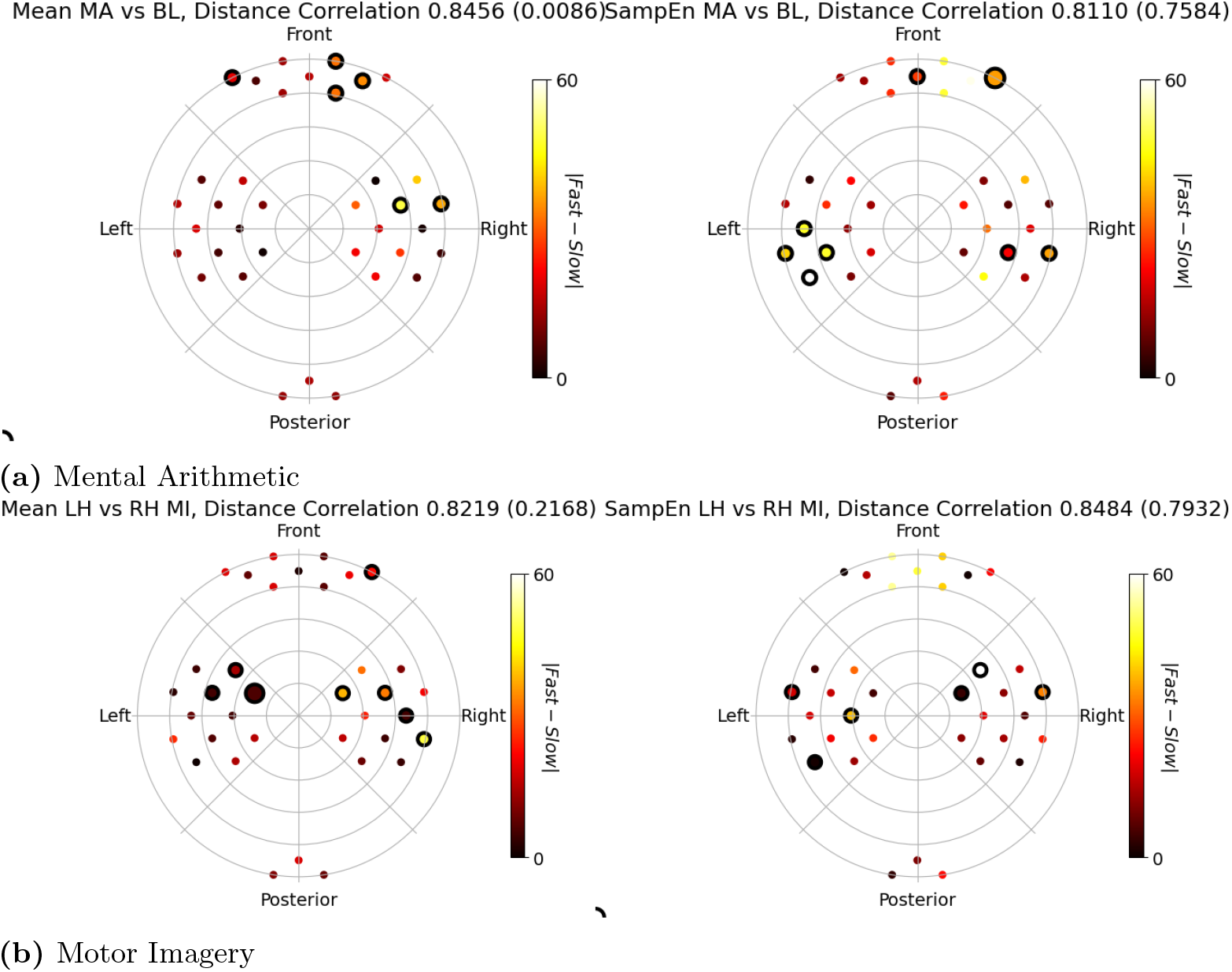
Results from comparing fast wave cardiac rhythms in fNIRS (0.8 Hz to 3 Hz) and traditional slow wave fNIRS (0 Hz to 0.2 Hz). Effect size displayed in the plots are the difference between the medians at the group level between fast and slow wave measures (either mean in the left column or sample entropy in the right column). Group analysis Wilcoxon signed rank tests were used on the group level to assess whether a detector center has a significantly different median for two tasks, where surviving detectors are demarcated by a thick marker outline. (a) displays mental arithmetic vs baseline results, while (b) displays results comparing left vs right hand motor imagery. The colors in the plots represent which modality (fast or slow wave) was significant for the given measure. In the title of each subplot, the distance correlation among the surviving detectors between fast and slow wave topoplots is provided, with the p-value in parenthesis.

## Conclusion

We conclude from this study that nonlinear analysis can detect different representations when using either fast (0.2 Hz to 0.6 Hz) or standard slow (< 0.2 Hz) wave frequencies in fNIRS data, which traditional methods like mean estimates can not. While origin of neural hemodynamic activity may be associated with oscillations at frequencies < 0.2 Hz, nonlinear interactions between such oscillations may indeed generate fNIRS oscillations at higher frequency. Thus, when assessing effects on fNIRS, comprehensive characterizations should also consider nonlinear properties of high frequency band oscillations. Furthermore, differences in the median in group analysis was found in different cortical regions depending on whether fast or slow wave fNIRS data was used with SampEn, which may reveal a modulation of systems that control arteriole vasomotor tone induced by nonlinear sympathovagal interactions, though further studies that engage or stimulate these systems need to be explored.

## Acknowledgments

The research leading to these results has received partial funding from the European Commission Horizon 2020 Program under grant agreement n. 813234 of the project RHUMBO and by the Italian Ministry of Education and Research (MIUR) in the framework of the CrossLab project (Departments of Excellence).

